# The Onset of Breathing at Birth Switches Off Regenerative Capacity

**DOI:** 10.64898/2026.10.03.756425

**Authors:** Xuan Song, Xinyi Cai, Shane V. Hegarty, Davide Speri, Masahiro Yasuda, Ryan J. Donahue, Joanna Stanicka, Ting Yan, Susu Mao, Sarah Yang, Jiawen Chen, Junjie Zhu, James K. Kiraly, Cheng-Hao Chien, Evgenii Kegeles, Daniel Feng, Kevin C. Stieger, Zhiyun Yang, Yiming Zhang, Gary Yellen, Hisashi Umemori, Philip R. Williams, Zhigang He

**Author notes:** Correspondence: Zhigang He. These authors contributed equally to this work (Xuan Song, Xinyi Cai, Shane V. Hegarty, Davide Speri).

## Abstract

Many mammalian tissues, including the central nervous system (CNS), lose regenerative capacity in adulthood1-3, but when and how this transition is initiated remains unclear. By monitoring mTOR activity, an evolutionarily conserved readout of cellular growth and intrinsic regenerative potential4-11, we found that mTOR signaling declines within minutes after birth in murine CNS neurons and in other organs such as the heart and skeletal muscles. Focusing on retinal ganglion cells (RGCs), we show that this abrupt decline is driven primarily by CO₂ exhalation following the onset of breathing after birth. Using real-time in vivo imaging of RGCs during the earliest postnatal period, we further discover that breathing-induced CO₂ exhalation rapidly causes intracellular alkalinization, activates an AMPK-family kinase, SNF-related kinase (SNRK)12-14, which in turn suppresses mTOR signaling. Importantly, this process remains reversible in adulthood: CO₂ inhalation or Snrk deletion restores mTOR activity in RGCs and enables robust regeneration of their axons after injury. Similar CO₂ inhalation–induced mTOR reactivation occurs across multiple cell types and organs, suggesting a general mechanism by which breathing-associated CO₂/pH signaling shapes the regenerative capacity across life and pointing to a potential CO₂-based strategy for promoting tissue regeneration.

---

Injured axons in the adult mammalian central nervous system (CNS) fail to regenerate, severely limiting recovery after trauma or neurodegenerative disease. Prior work has identified multiple molecular regulators of axon growth^9,15–17^. Among them, mTOR has emerged as a key regulator of neuronal intrinsic regenerative capacity across species, in part through its role in activating anabolic metabolism and supporting the production of cellular building blocks for axon regrowth^4–11^. However, a fundamental question remains unresolved: when, how, and why do mammals lose intrinsic regenerative capacity during development? Defining the biological events that terminate neuronal growth competence may reveal strategies to restore regenerative ability in the adult CNS and other tissues.

Evidence suggests that regenerative capacity declines in multiple systems during early postnatal life. Neonatal mice can mount scar-free wound healing responses that support axon growth through the injury site, yet this capacity disappears within the first postnatal week^18^. Similarly, dissociated retinal ganglion cells (RGCs) from embryonic rodents exhibit robust neurite outgrowth, whereas postnatal RGCs show markedly reduced growth competence^19^. Consistently, the activity of mTOR is dramatically reduced in postnatal RGCs^4^. Furthermore, comparable developmental declines occur in other tissues, including the heart, skin, and sensory hair cells^20–26^. These observations raise the possibility that a shared physiological transition in early life actively restricts regenerative potential across tissues in mammals.

Birth represents one of the most dramatic and critical physiological transitions in mammalian life. Within minutes, the newborn shifts from a protected intrauterine environment to independent homeostatic regulation^27–29^. This transition requires coordinated adaptations, including the onset of pulmonary respiration^29–32^, closure of fetal circulatory shunts^33–35^, transition from placental nutrient supply to feeding^36–39^, and initiation of thermoregulation^27,40,41^. Although these events enable survival outside the uterus, whether they also contribute to the loss of regenerative capacity is unknown. In principle, regenerative ability could decline gradually through intrinsic developmental programs. However, a discrete physiological event associated with birth could also trigger a rapid shutdown of growth-promoting pathways. RGCs provide a powerful system to test this possibility because of their experimental accessibility and their well-established axotomy models for studying CNS regeneration^17,42,43^.

Here, we show that the onset of respiration at birth rapidly suppresses intrinsic growth signaling through a CO₂/pH–SNRK–mTOR pathway. Monitoring mTOR activity revealed near-complete inactivation within minutes of birth in RGCs and multiple organs. Remarkably, this birth-associated mTOR suppression depends on SNRK, a noncanonical kinase of the AMPK family. Using transpupillary two-photon imaging *in vivo*, we found that the onset of breathing drives rapid CO₂ exhalation-dependent intracellular alkalinization in RGCs, followed by activation of SNRK and consequent suppression of mTOR signaling. Importantly, this pathway is reversible in mature neurons: either SNRK loss or elevated CO₂ exposure, which lowered intracellular pH, reactivates mTOR, and enables robust RGC axon regeneration after injury. Together, these findings identify respiration onset at birth as a major physiological switch that acutely restricts neuronal growth competence and establish a targetable CO₂/pH–SNRK–mTOR signaling axis linking postnatal respiration to the developmental loss of regenerative capacity in the adult CNS and possibly other tissues.

## mTOR is inactivated within minutes after birth

Previous studies have shown that mTOR activity, a robust correlate of neuronal regenerative capacity^4–8,10,11^, declines markedly between embryonic and postnatal stages^4^, but the precise timing and trigger of this decline have remained unclear. We first measured mTOR activity during early postnatal stages by examining phosphorylation of S6 (p-S6), a well-established downstream readout of mTOR signaling, in RGCs. As expected, before birth, most RGCs (RBPMS positive) were p-S6 positive (Fig. 1a, d). Strikingly, within 5 min of Caesarean section delivery, p-S6 signal in RGCs dropped sharply and was nearly eliminated (Fig. 1b, d). This birth-induced suppression of mTOR activity persisted for at least 12 hr after delivery (Fig. 1c, d), identifying birth as a critical developmental switch that rapidly alters the intrinsic growth state of RGCs.

**Figure 1.**
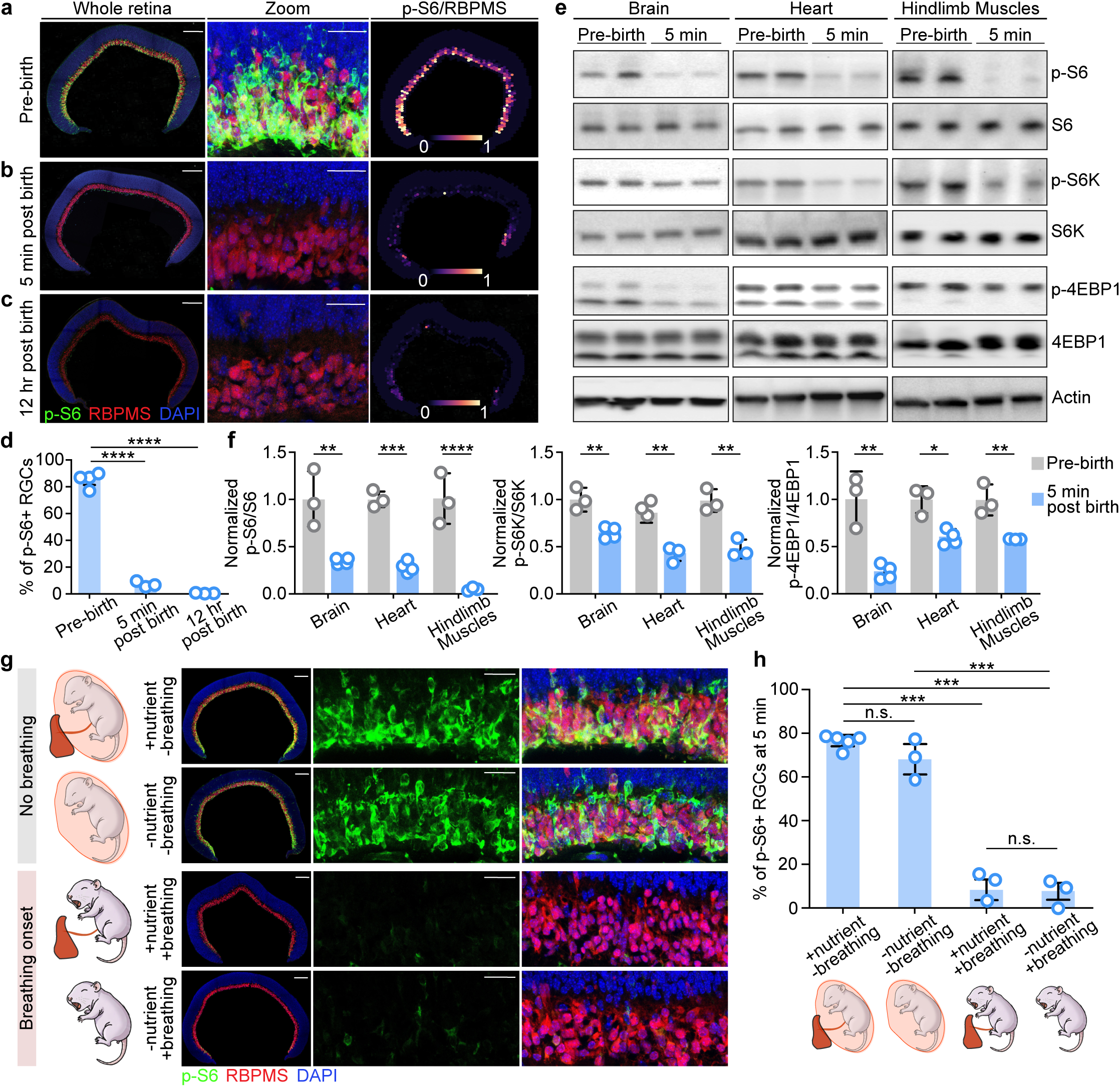
Breathing drives mTOR inactivation within minutes after birth. **a-c,** Representative images of phosphorylated S6 (p-S6, green), RBPMS (red), and DAPI (blue) staining in whole retinal sections (left), with magnified views (middle), and normalization of p-S6/RBPMS intensity (right) from pre-birth (E18.5) embryos, 5 min, and 12 hr postnatal pups. Scale bars, 200 μm (whole retinal section); 50 μm (magnified views). **d,** Quantification showing that the percentage of p-S6-positive RGCs significantly decreases after birth. Each circle represents the mean percentage of p-S6-positive RGCs among total RBPMS-positive RGCs in the ganglion cell layer (GCL) of each retina. For each whole retina section, all RGCs were included in the quantification. ****p < 0.0001; N = 4, 3 and 3 biological replicates for pre-birth, 5 min post-birth and 12 hr post-birth groups, respectively; unpaired Welch’s t-test. **e,** Western blot showing that mTOR activity markers (p-S6, p-S6K, and p-4EBP1) decrease markedly in brain, heart, and hindlimb muscles within 5 min after birth. Quadriceps femoris muscles and biceps femoris muscles from both left and right hindlimbs were pooled together to generate hindlimb muscles sample for each mouse. **f,** Quantification of Western blot result shown in **(e)**. *p < 0.05, **p < 0.01, ***p < 0.001; N = 3 and 4 biological replicates for pre-birth and 5 min post birth, respectively; unpaired Welch’s t-test. **g,** Experimental scheme (left) illustrating conditions with deprivation of maternal nutrient supply or inhibition of breathing onset. Representative images (right) of p-S6, RBPMS, and DAPI staining in whole retinal sections (from newborns at 5 min after birth) and corresponding magnified views under the indicated conditions. 200 μm (whole retinal section); 50 μm (magnified views). **h,** Quantification showing the percentage of p-S6-positive RGCs under different conditions. At 5 min after birth, deprivation of maternal nutrient supply does not alter p-S6 levels, whereas initiation of breathing significantly decreases mTOR activity. ***p < 0.001; n.s., not significant; N = 5, 3, 3 and 3 biological replicates for (“+ nutrient - breathing”), (“- nutrient - breathing”), (“+ nutrient + breathing”) and (“- nutrient + breathing”) groups, respectively; unpaired Welch’s t-test.

We next asked whether this response is specific to RGCs or reflects a broader systemic transition. To address this, we examined p-S6 and additional mTOR pathway markers by western blotting and immunofluorescence in other tissues, including brain, heart, and hindlimb skeletal muscles. We observed similarly rapid mTOR inactivation across multiple cell types and tissues (Fig. 1e, f, and Extended Data Fig. 1), indicating that birth-associated mTOR suppression is a widespread, cross-body event. Importantly, phosphorylation of multiple downstream mTOR targets, including S6, S6K, and 4EBP1, was reduced after birth, whereas total protein levels remained unchanged (Fig. 1e). These findings argue against a mechanism driven by slower transcriptional or translational changes and instead support the idea that birth triggers an acute regulatory signal that rapidly suppresses mTOR activity.

## Breathing drives rapid mTOR inactivation at birth

Birth imposes abrupt physiological changes, including loss of umbilical nutrient supply, initiation of pulmonary respiration, and exposure to ambient temperature. We therefore asked if any of these transitions drives the immediate mTOR inactivation observed at birth. A shift in body temperature was not sufficient: maintaining pups at 37 °C immediately after Caesarean delivery failed to preserve mTOR activity (data not shown).

We next considered two concurrent birth-associated events: cessation of maternal nutrient delivery and onset of breathing^33–39,44^. In the Caesarean delivery-associated perinatal setting, these variables can be experimentally dissociated, as the umbilical cord sustains maternal nutrient transfer, whereas the intact amniotic sac prevents air breathing. We therefore examined mTOR activity in RGCs of newborn pups with or without an intact umbilical cord, and either within or outside the amniotic sac.

Preservation of maternal nutrient supply alone did not prevent the loss of p-S6 in RGCs 5 min after Caesarean delivery (“+ nutrient + breathing” and “− nutrient + breathing”) (Fig. 1g, h). Consistent with this, blood glucose was not reduced for at least 5 min after birth (Extended Data Fig. 2a), despite near-complete suppression of mTOR signaling during this interval. Thus, acute nutrient withdrawal is insufficient to account for the minute-scale, rapid loss of mTOR activity after birth. Although postnatal feeding restored mTOR activity in a subset of RGCs (Extended Data Fig. 2b, c), consistent with prior evidence that only a minority of mature RGCs are p-S6 positive ^4,45^, this response did not explain the rapid suppression observed at delivery.

By contrast, the pups that remained enclosed within the amniotic sac (“− nutrient − breathing”) retained markedly higher mTOR activity than air-exposed pups for at least 5 min and up to 30 min after delivery (Fig. 1g, h and Extended Data Fig. 2d, e). Conversely, nearly all RGCs became p-S6-negative in pups exposed to air and breathing, both with (“+ nutrient + breathing”) or without (“− nutrient + breathing”) intact umbilical cord (Fig. 1g, h). The decline in p-S6 signal started within 4 min after breathing onset and was nearly complete after 5 min (Extended Data Fig. 2f). Thus, birth-associated mTOR inactivation is not driven simply by loss of maternal nutrient input but is tightly linked to the sudden transition into the air-breathing state for the first time at birth.

We then asked what upstream mechanisms could directly inhibit mTOR activity following breathing onset. Two canonical mTOR regulators are the growth factor-sensitive tuberous sclerosis complex (TSC) pathway and the amino acid-dependent Rag pathway, which repress or activate mTOR, respectively ^10,11^. To assess their contributions to birth-induced mTOR inactivation, we knocked out *Tsc1* in RGCs and measured their p-S6 signal across birth. Unlike wild-type RGCs, TSC-deficient RGCs from *Tsc1 ^fl/fl^; Vglut2-Cre* conditional knockout (cKO) mutants significantly maintained p-S6 levels across birth (Extended Data Fig. 2g, h), demonstrating the necessity of intact TSC activity for post-birth mTOR inactivation. Thus, these observations demonstrate that birth-associated mTOR inactivation can be explained by upstream TSC-induced repression, likely triggered by the acute onset of autonomous respiration at birth.

## SNRK, a non-canonical AMPK-like kinase, is required for breathing-induced rapid mTOR inactivation in RGCs

How might respiration at birth repress mTOR activity? We hypothesized that AMPK-family kinases (AMPK-F) signaling might contribute to this for at least two reasons. First, AMPK-F kinases are conserved stress sensors that inhibit mTOR^46–48^, and one of the mediators is the TSC^49–51^, making them plausible links between birth-induced stress and mTOR repression. Second, we performed an independent CRISPR-based kinase screen for regulators of axon regeneration after adult optic nerve injury (Extended Data Fig. 3a) and identified SNRK, an AMPK-F member^12–14^, as the strongest negative regulator of axon regeneration (Extended Data Fig. 3b-d). Consistently, conditional deletion of *Snrk* in RGCs by intravitreal AAV2-Cre delivery under the RGC-selective mSncg promoter^52^ in *Snrk ^fl/fl^* mice at P28 induced robust mTOR activation 14 days after injection (Fig. 2a, b) and enhanced axon regeneration 14 days post-injury (Fig. 2c-e). Together, these results support a hypothesis in which birth-associated respiration activates AMPK-F signaling, including SNRK, to repress mTOR and restrict the regenerative state of RGCs.

**Figure 2.**
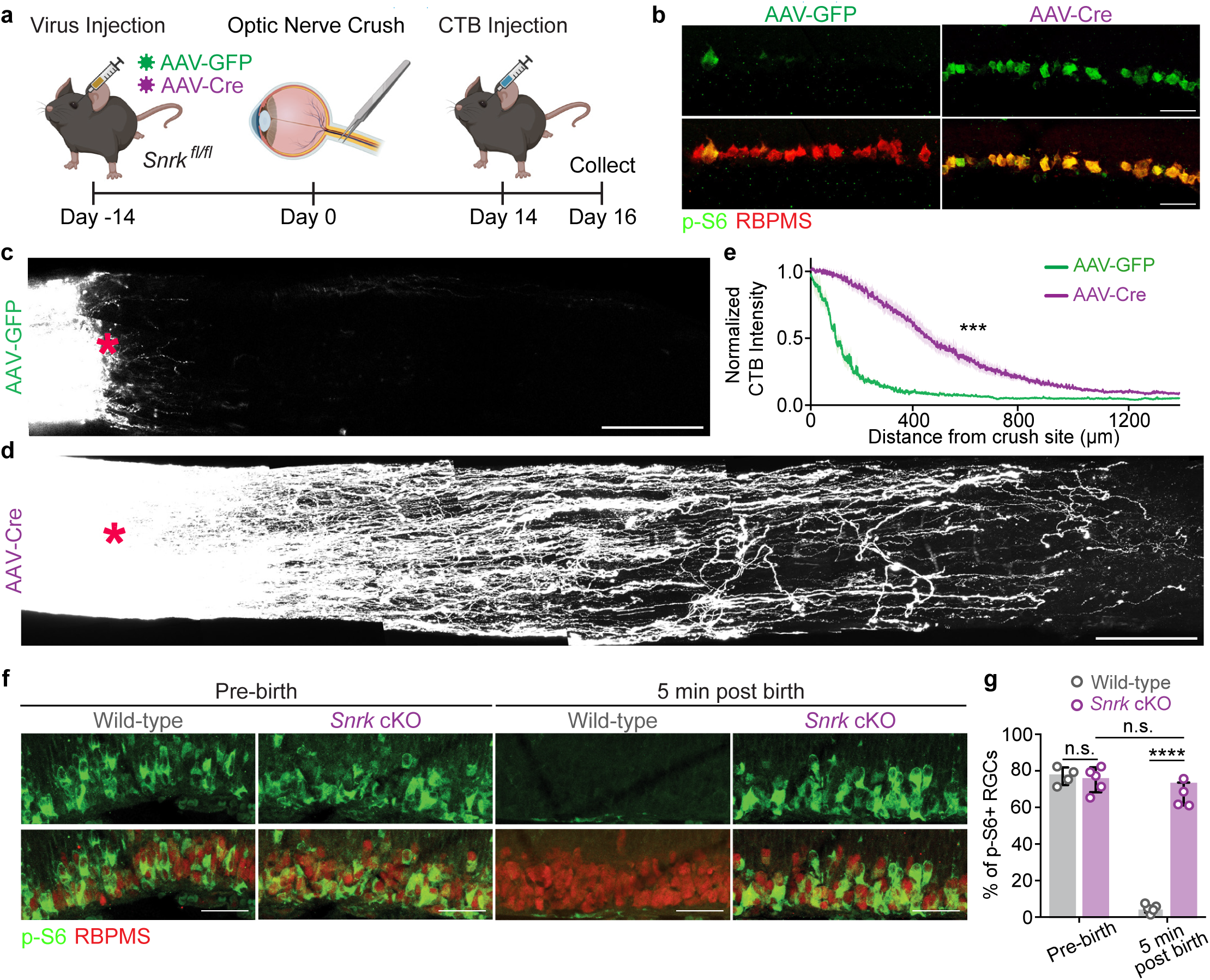
SNRK is required for breathing-induced rapid mTOR inactivation in RGCs. **a,**Experimental scheme for intravitreal injection of AAV2-mSncg-Cre or GFP control virus and optic nerve crush in *Snrk ^fl/fl^* mice. **b,** Representative images of phosphorylated S6 (p-S6, green) and RBPMS (red) staining in the retina section of *Snrk ^fl/fl^* mice injected with either GFP or Cre-expressing AAVs. Scale bar, 20 µm. **c, d,** Representative full thickness optic nerve images showing CTB-labeled axons at 14 days after injury in *Snrk ^fl/fl^* mice injected with AAV-GFP **(c)** or AAV-Cre **(d)**. Scale bar, 100 µm. The red asterisk indicates the ONC site. **e,** Quantification of CTB-labeled fluorescent intensity distal to the crush site in *Snrk ^fl/fl^* mice injected with AAV-Cre or AAV-GFP. Data are shown as mean ± SEM; N = 7 and 8 biological replicates for AAV-GFP and AAV-Cre injected mice, respectively. ***p < 0.001; mixed-effects analysis. **f,** Representative images of phosphorylated S6 (p-S6, green) and RBPMS (red) staining in the retinal section of pre-birth (E18.5) embryos and 5 min post-birth newborns of wild-type or *Snrk* cKO. Scale bar, 20 µm. **g,** Quantification showing the percentage of p-S6-positive RGCs. Data are presented as mean percentages of p-S6-positive and RBPMS-positive cells among total RBPMS-positive cells in the GCL of each retina. ***p < 0.001; n.s., not significant; N = 4 wild-type and 5 *Snrk* cKO biological replicates for pre-birth; N = 5 wild-type and 4 *Snrk* cKO biological replicates at 5 min post birth; unpaired Welch’s t-test.

Notably, all 13 AMPK-F members were included in our gRNA library; however, SNRK was the only member that met the inclusion criteria (axons regenerated ≥ 500 μm) in the initial forward genetic screen (see Methods). One possible explanation is the differential developmental expression of these AMPK-F members. For example, Mark1/2/3 are highly expressed in embryonic RGCs (Extended Data Fig. 3e) and have established roles in neuronal polarity during differentiation^53,54^. In contrast, *Snrk* remains among the most highly expressed AMPK-F kinases in RGCs from late embryonic stages through adulthood (Extended Data Fig. 3e). This expression pattern prompted us to test whether SNRK contributes to birth-induced mTOR inactivation. To do so, we selectively deleted *Snrk* in embryonic RGCs using *Snrk ^fl/fl^; Vglut2-Cre* mice and measured p-S6 levels across birth. Strikingly, SNRK-deficient RGCs retained a strong p-S6 signal at 5 min after birth, at levels comparable to those in embryonic RGCs (Fig. 2f, g). These results demonstrate that SNRK-dependent AMPK-F signaling links respiration-driven birth stress to the rapid suppression of mTOR activity, and thereby regenerative capacity.

## SNRK predominantly contributes to the immediate AMPK activation following birth

Next, we asked whether activation of SNRK accompanies respiration-associated suppression of mTOR and, if so, whether SNRK activation precedes mTOR activity suppression. Because no selective *in vivo* readout for SNRK activity is currently available, we monitored total AMPK-F activity in RGCs and then determined the contribution of SNRK to that signal. Because mTOR activity is silenced within minutes after breathing onset, the relevant upstream kinase activity must be measured continuously in real time. We therefore used the ratiometric reporter ExRai-AMPKAR^55^, which contains the MRRVATLVD phosphoacceptor motif and reports the activity of AMPK-F, including SNRK^56^. Upon phosphorylation, the MRRVATLVD motif binds to the phosphoamino acid-binding forkhead associated domain 1 (FHA1), inducing a conformational change in the sensor that increases the emission ratio upon 920/800 nm excitation^55^.

To record changes in AMPK-F activity at birth, we adapted transpupillary two-photon imaging for newborn mice (Fig. 3a and Extended Data Fig. 4-9; Supplementary Video 1; see Methods). ExRai-AMPKAR was sparsely expressed in RGCs by intravitreal delivery of AAV/DJ-CAG-DIO-ExRai-AMPKAR to E13 *Vglut2-Cre* embryos^57^, a strategy that enabled reliable sensor expression and single-cell segmentation at the time of imaging and data analysis (Extended Data Fig. 5). To preserve physiological birth timing while allowing immediate optical access, pups were delivered by Caesarean section at E18.5 and imaged directly in a custom holder (Fig. 3a and Extended Data Fig. 4b). Because pups delivered by Caesarean section require postnatal tactile stimulation to initiate respiration, residual amniotic fluid was cleared and tactile stimulation was applied in a standardized manner (Extended Data Fig. 6a, b; Supplementary Video 2). Upon stimulation, newborn mice passed through an initial gasping phase followed by eupnea (Extended Data Fig. 6c, d), with breathing frequency rising from 7 ± 4 breaths per min (n = 38) at 1 min to 45 ± 7 breaths per min (n = 33) at 5 min. Imaging and analysis conditions were optimized to maximize ratiometric stability during longitudinal acquisition, including calibration of excitation power, restriction of run length, focal-plane selection, motion correction, and channel registration (Extended Data Fig. 7-9; see Methods).

**Figure 3.**
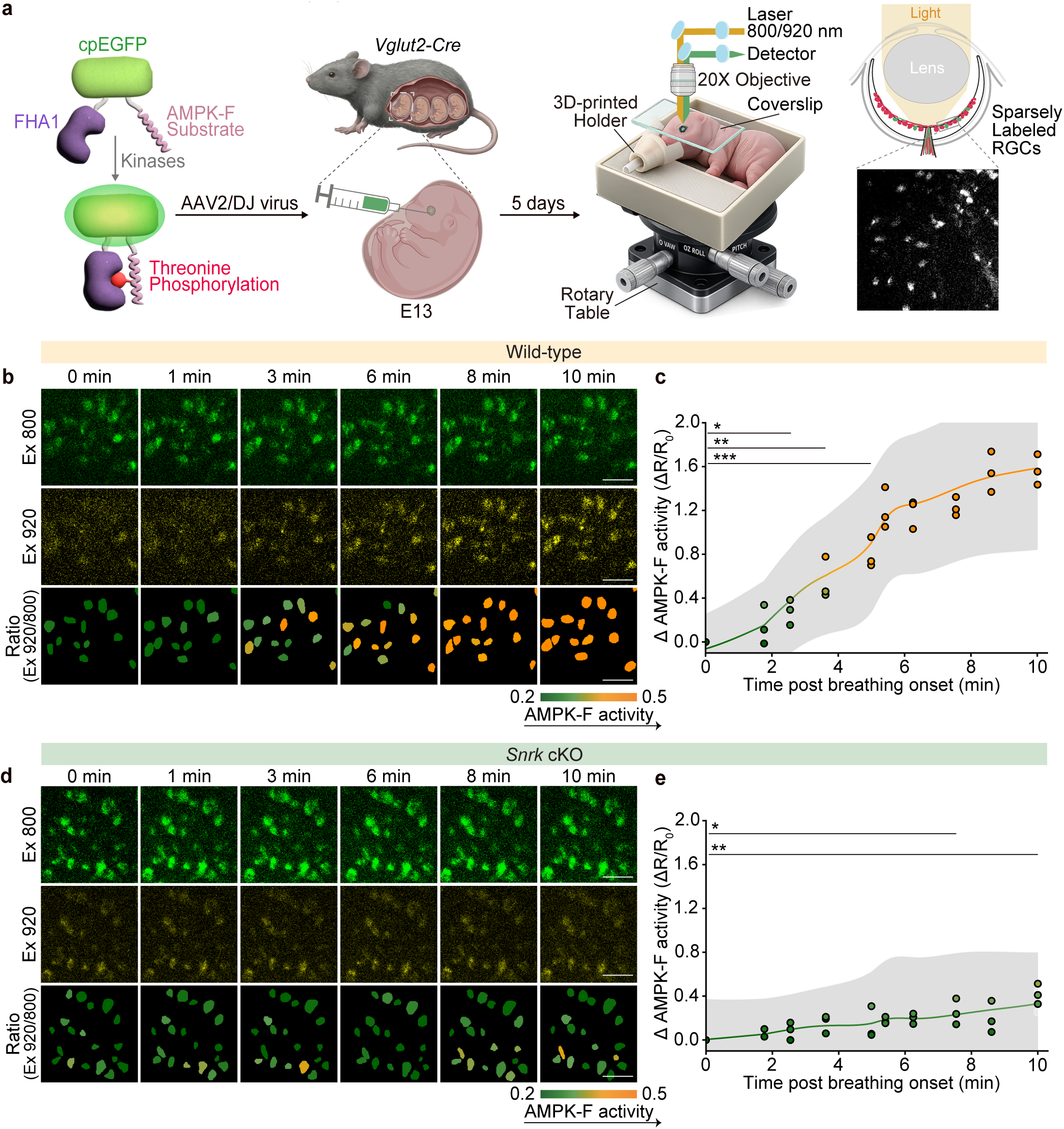
SNRK predominantly contributes to the rapid AMPK activation following birth. **a,**Experimental scheme illustrating the ExRai-AMPKAR sensor expression via *in utero* injection and the *in vivo* two-photon imaging setup of neonatal mice. **b,** Representative images of ExRai-AMPKAR in *Vglut2-Cre* pups acquired at 800 nm and 920 nm excitation wavelengths at the indicated time after breathing onset. **c,** Quantification of the emission ratio (920/800 nm excitation) of ExRai-AMPKAR in *Vglut2-Cre* pups after breathing onset. Raw ratio from the post-breathing stage was normalized to the value of first frame to visualize the signal changes. Data are shown as mean ± SD; N = 3 biological replicates; n = 97 RGCs. *p < 0.05, **p < 0.01, ***p < 0.001; mixed-effects analysis with FDR correction. **d,** Representative images of ExRai-AMPKAR in *Vglut2-Cre; Snrk^fl/fl^* (*Snrk* cKO) pups acquired at 800 nm and 920 nm excitation wavelengths. **e,** Quantification of the emission ratio (920/800 nm excitation) of ExRai-AMPKAR in *Snrk* cKO pups after breathing onset. Raw ratio from the post-breathing stage was normalized to the value of the first frame to visualize the signal changes. Data are shown as mean ± SD; N = 3 biological replicates; n = 168 RGCs. *p < 0.05, **p < 0.01; mixed-effects analysis with FDR correction.

First, we validated the reporter *in vivo*. Intravitreal delivery of rotenone plus oligomycin, which imposes acute energy depletion^58^, rapidly increased ExRai-AMPKAR signal in newborn RGCs (Extended Data Fig. 10a). 5-Aminoimidazole-4-carboxamide-ribonucleoside (AICAR), an AMPK-F agonist^59^, likewise induced a rapid increase, followed by partial recovery (Extended Data Fig. 10b). Ionomycin, which activates AMPK-F through calcium/CaMKK signaling^60^, elevated the sensor activity within 2 min (Extended Data Fig. 10c). These responses are consistent with the known sensitivity of AMPK-F to energetic stress and calcium-dependent signaling^61–63^.

Using this imaging paradigm, we monitored AMPK-F kinase activity in the same RGCs before and after breathing onset. ExRai-AMPKAR revealed a rapid increase in AMPK-F activity after breathing onset, measured as an increase in the emission ratio from 920/800 nm excitation (Fig. 3b, c). This response was not observed with a phospho-dead reporter carrying a Thr-to-Ala substitution in the target motif (Extended Data Fig. 11), indicating that the signal increase might reflect kinase-dependent reporter phosphorylation. The initial increase of AMPK-F activity occurs 2-3 min after breathing onset and continues thereafter (Fig. 3c), temporally preceding but overlapping with mTOR inactivation. AMPK-F activity continued to increase until it stabilized approximately 8 min after breathing onset (Fig. 3c). However, in pups that did not initiate breathing, reporter activity remained largely unchanged (Extended Data Fig. 12a). Thus, activation of AMPK-F signaling is tightly coupled to the onset and progression of respiration.

Because ExRai-AMPKAR reports the integrated activity of AMPK-F, we next asked what proportion of the postnatal AMPK-F signal activation could be attributed to SNRK. Using the same strategy, we expressed ExRai-AMPKAR in RGCs of E13 *Snrk ^fl/fl^; Vglut2-Cre* embryos and monitored the reporter activity across birth. Loss of SNRK markedly attenuated the postnatal rise in ExRai-AMPKAR signal (Fig. 3d, e): the initial increase was largely abolished, and the later phase was strongly diminished, whereas non-breathing newborns remained at baseline level throughout (Extended Data Fig. 12b). These results indicate that the respiration-triggered increase in AMPK-F activity in newborn RGCs is mediated predominantly through SNRK activation. Although biosensor output can be influenced by environmental variables such as pH and temperature ^64^, the direct comparison of control and SNRK-deficient RGCs under the same perinatal conditions argues that the genotype-dependent difference in signal primarily reflects loss of SNRK-dependent phosphorylation of the reporter rather than nonspecific environmental effects. On the other hand, the residual signal in SNRK-deficient RGCs (Fig. 3e) may arise from incomplete elimination of AMPK-F input, reflecting activity of other intact family members, and/or from environmental modulation of sensor behavior. These findings support a model in which the onset of respiration activates SNRK, which in turn accounts for the majority of the AMPK-F activity detected by ExRai-AMPKAR at birth.

Together, these data place SNRK activation upstream of birth-associated mTOR shutdown. Respiration onset triggers a rapid, predominantly SNRK-dependent AMPK-F response in RGCs, and this response precedes the loss of mTOR activity, consistent with a causal role in coupling the acute transition of air breathing and the rapid postnatal mTOR repression.

## Breathing-initiated CO₂ exhalation contributes to both SNRK activation and mTOR inactivation

What accounts for the minute-scale activation of AMPK-F and suppression of mTOR signaling pathways at birth? A primary physiological consequence of respiration is a simultaneous rise in systemic O₂ and drop in CO₂. To distinguish between these two variables, we delivered pups into controlled gas environments during imaging, selectively altering either CO₂ or O₂ levels (Fig. 4a, c). Lowering oxygen did not attenuate the increase in AMPK-F activity during the first 8-9 min after birth (Fig. 4c, d), arguing that the acute response is not driven primarily by increased O₂ availability. In contrast, elevating CO₂ at birth effectively blocked AMPK-F activation over the first 10 min (Fig. 4a, b). The same manipulation affected mTOR activity as well. Measurement of p-S6 showed that 10% CO₂, but not 10% O₂, prevented the postnatal loss of mTOR activity seen under ambient air conditions (Fig. 4e, f). Together, these results identify CO₂ clearance, rather than oxygenation, as the critical respiratory signal that couples breathing onset to rapid SNRK/AMPK-F activation and concomitant mTOR inactivation at birth.

**Figure 4.**
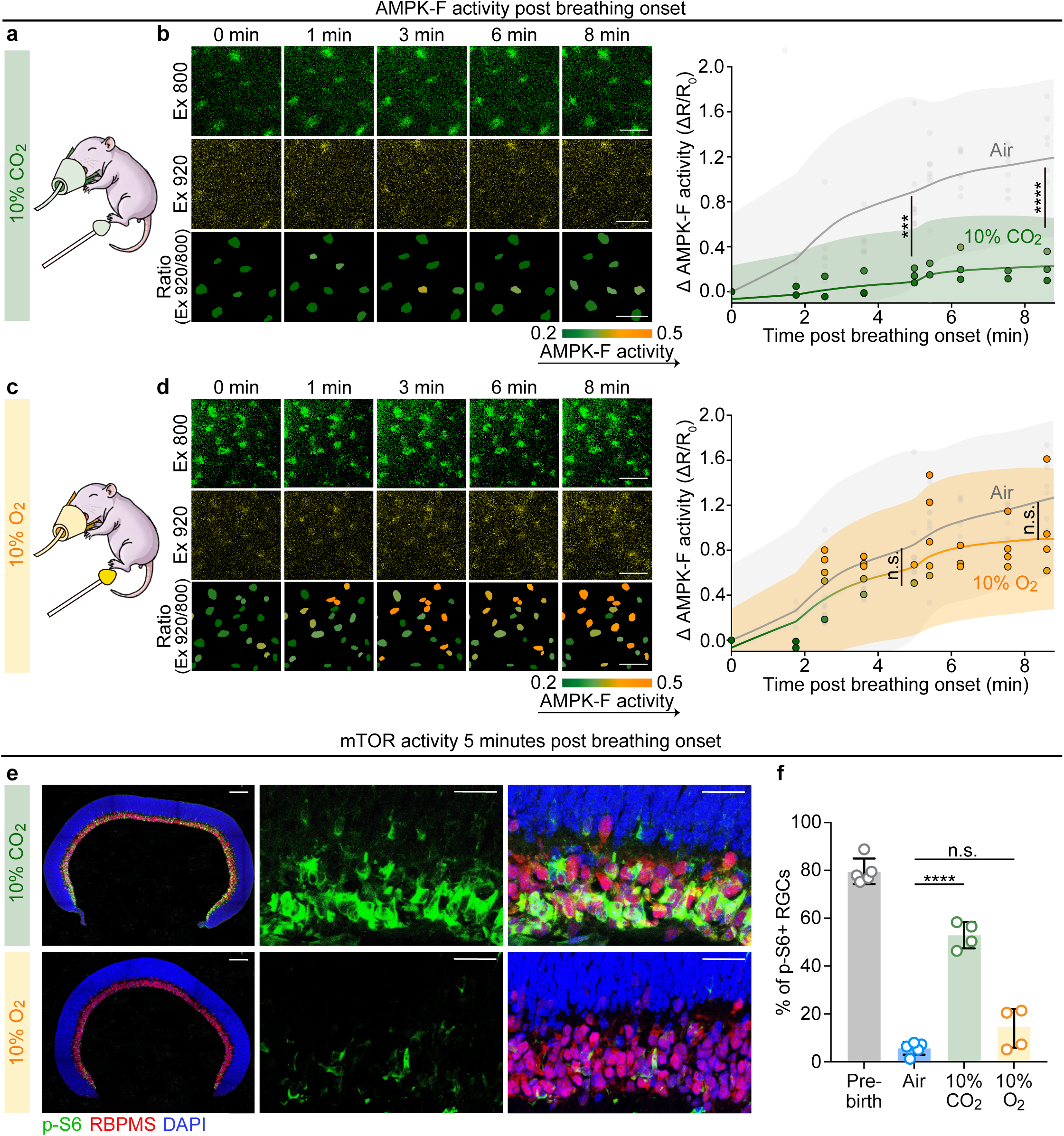
Breathing-initiated CO_2_ exhalation significantly contributes to both SNRK/AMPK activation and mTOR inactivation. **a,**Experimental scheme of newborns delivered under elevated 10% CO_2_ conditions for *in vivo* imaging of AMPK-F activity. **b,** Representative images (left) of ExRai-AMPKAR in *Vglut2-Cre* pups acquired at 800 nm and 920 nm excitation wavelengths. Quantification (right) showing the emission ratio (920/800 nm excitation) of ExRai-AMPKAR. Raw ratio from the post-breathing stage was normalized to the value of the first frame to visualize the signal changes. Data are shown as mean ± SD; N = 3 biological replicates; n = 95 RGCs. ***p < 0.001, ****p < 0.0001; mixed-effects analysis with FDR correction. **c,** Experimental scheme of newborns delivered under reduced 10% O₂ conditions for *in vivo* imaging of AMPK-F activity. **d,** Representative images (left) of ExRai-AMPKAR in *Vglut2-Cre* pups acquired at 800 nm and 920 nm excitation wavelengths. Quantification (right) showing the emission ratio (920/800 nm excitation) of ExRai-AMPKAR. Raw ratio from the post-breathing stage was normalized to the value of the first frame to visualize the signal changes. Data are shown as mean ± SD; N = 5 biological replicates; n = 88 RGCs; n.s., not significant; mixed-effects analysis with FDR correction. **e,** Representative images of phosphorylated S6 (p-S6, green), RBPMS (red), and DAPI (blue) staining in retinal sections from 5 min postnatal pups delivered in 10% CO_2_ (upper row) and 10% O_2_ (bottom row). Scale bars, 200 μm (whole retina) and 50 μm (magnified views). **f,** Quantification showing that postnatal exposure to 10% CO_2_, but not 10% O_2_, prevents the reduction in p-S6-positive RGCs in newborns 5 min after breathing onset. Data are presented as the mean percentages of p-S6-positive cells among total RGCs in the ganglion cell layer (GCL) of each retina. ****p < 0.0001; n.s., not significant; N = 5 biological replicates for the pre-birth group and N = 5, 4 and 4 biological replicates for pups born in air, 10% CO₂ and 10% O₂ for 5 min, respectively; unpaired Welch’s t-test.

## Breathing-triggered intracellular alkalinization activates SNRK

Next, we asked how breathing activates SNRK in newborn RGCs. AMPK-F kinases are classically engaged by either energetic stress or calcium/CaMKK signaling^61–63^, so we used our *in vivo* imaging pipeline to test both possibilities directly in RGCs from newborns. ATP was monitored with ATeam1.03^65,66^ (Extended Data Fig. 13a, b), and calcium with Twitch-2B^67,68^ (Extended Data Fig. 13c-f). Cells were segmented using a U-net-based network (Extended Data Fig. 14; see Methods). FRET efficiency was quantified after correction for bleed-through (Extended Data Fig. 15c; see Methods) and normalization to photon flux following the previously established methods^69^ (see Methods). ATP levels remained stable throughout the first 10 min after birth, including the period when AMPK-F activity was already rising (Extended Data Fig. 13a, b), arguing against acute energy depletion as the trigger. This interpretation is further supported by the fact that SNRK, unlike canonical AMPK-F complexes, lacks the AMPK γ subunit that confers sensitivity to AMP/ATP ratios^70^. Thus, the early AMPK-F response is unlikely to be driven primarily by energetic stress.

Intracellular calcium concentrations, as indicated by Twitch-2B signal, remained stable for the first 4 min after breathing onset, then increased sharply afterwards (Extended Data Fig. 13c, d), and plateaued near 6 min, whereas calcium remained stable in non-breathing newborns after birth (Extended Data Fig. 13e). Pups delivered into 10% CO₂ also showed lower postnatal calcium levels than littermates in ambient air (Extended Data Fig. 13f). Because this calcium rise occurred after the initial increase in SNRK/AMPK-F activity, calcium-dependent signaling is unlikely to account for the initial kinase activation. However, the increase in calcium levels may contribute to the sustained rise in AMPK-F activity during the later phase. Thus, these results suggest that the acute SNRK/AMPK-F activation in newborn RGCs may result from a third factor.

One attractive candidate for a third respiration-linked variable is CO₂-dependent pH regulation, as Snf1, the yeast homolog of mammalian AMPK-F, can be activated by different environmental stressors, including alkaline pH ^71^. Within the first minutes after breathing onset, CO₂ exhalation shifts the CO₂/HCO₃⁻/H⁺ equilibrium toward lower proton concentration, predicting rapid alkalinization in blood. However, it is unknown whether and how fast this will impact the intracellular pH (pHi) of RGCs. To test this, we monitored pHi *in vivo* using the FRET-based sensor pHLemon-rev, a configuration optimized for sensitivity in the physiological range of pH 7-8 ^72^. It contains a pH-insensitive mTurquoise2 (as donor) and a pH-sensitive EYFP (as acceptor) (Extended Data Fig. 15d). As intracellular pH increases, energy transfer efficiency from the pH-insensitive donor fluorophore mTurquoise2 to the pH-sensitive acceptor fluorophore EYFP is expected to increase^72^. The sensor was expressed in RGCs by intravitreal delivery of AAV/DJ-CAG-DIO-pHLemon-rev into E13 *Vglut2-Cre* embryos, and EYFP/mTurquoise2 signals were imaged simultaneously under 820 nm excitation at birth with low laser intensity to avoid tissue damage (Extended Data Fig. 16a) during longitudinal imaging. Cells were segmented using the same U-net-based network as mentioned above (Extended Data Fig. 14; see Methods). Bleed-through was calculated across different pH conditions as described above (Extended Data Fig. 15c and e; see Methods). FRET efficiency was quantified after correction for bleed-through and normalization to photon flux as described above (Extended Data Fig. 15e; see Methods). pH-associated FRET difference was not sensitive to temperature changes from 25°C to 45 °C (Extended Data Fig. 16b). Predicted pH using FRET efficiency calculation in HEK293T cells closely matched the pre-determined pH values, validating our FRET imaging and calculation system (Extended Data Fig. 15f-g).

pHi in RGCs rose rapidly after breathing onset. FRET efficiency increased within ∼1 min and continued to rise until stabilizing after 5 min (Fig. 5a, b), indicating rapid intracellular alkalinization in RGCs. No such change was observed in the RGCs of the pups that did not initiate breathing (Fig. 5c, d). Moreover, when pups were delivered into 10% CO₂, this alkalinization was blocked: pHi remained low and further decreased to below embryonic levels when 10% CO₂ was prolonged (Extended Data Fig. 17) whereas littermates in ambient air underwent the expected rise in pHi (Fig. 5a, b). Thus, CO₂ clearance is both necessary and temporally coupled to the rapid intracellular alkalinization that follows birth.

**Figure 5.**
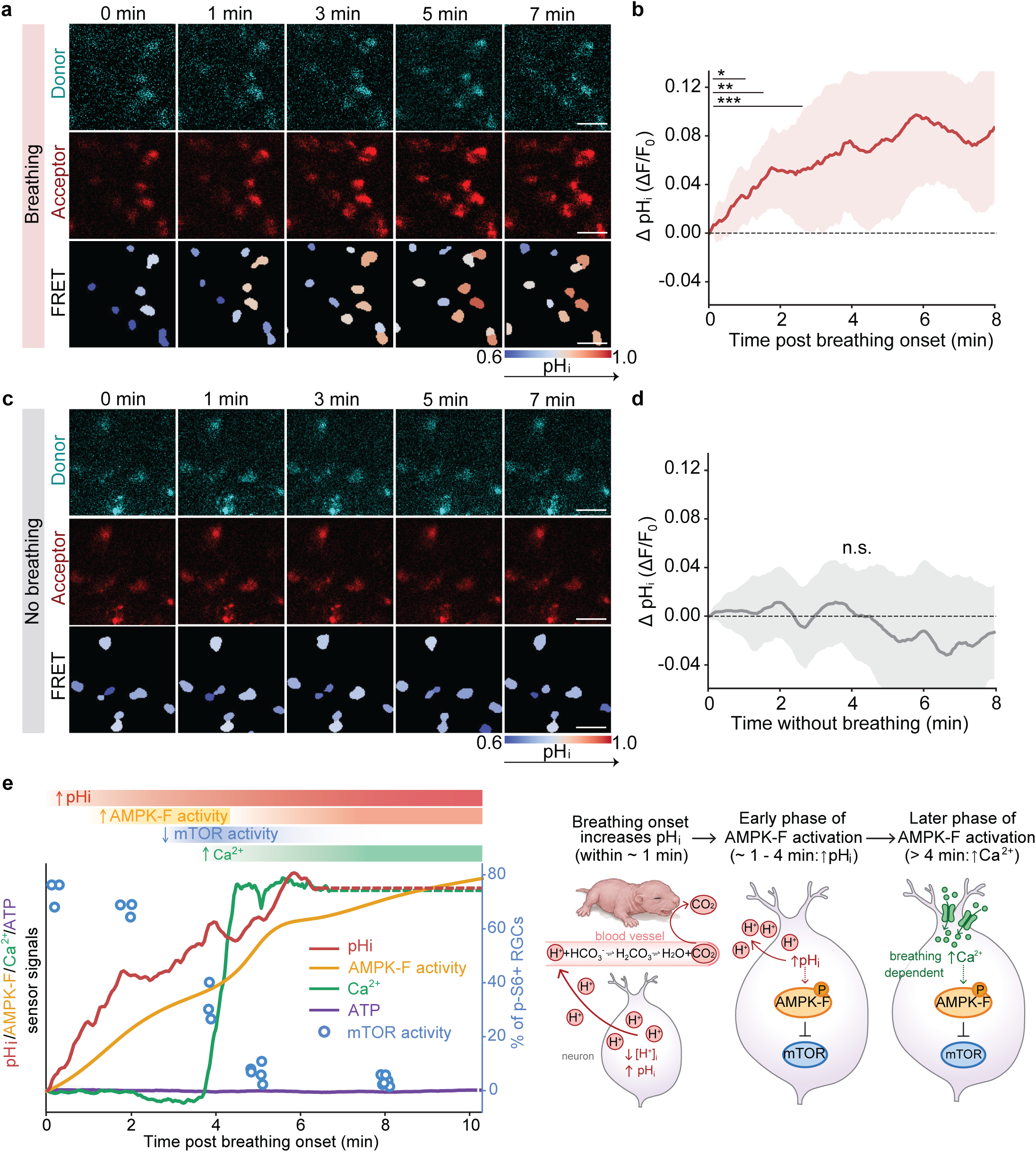
Breathing triggers intracellular alkalinization. **a,**Representative images of donor channel (upper row), acceptor channel (middle row), and calculated FRET efficiency (bottom row) of pHLemonrev in *Vglut2-Cre* RGCs from newborns acquired with 820 nm excitation after breathing onset. **b,** Quantification showing increased FRET efficiency of pHLemon-rev in RGCs from newborns after breathing onset. Raw FRET values from the post-breathing stage were normalized to the value of the first frame to visualize the signal changes. Data are shown as mean ± SD; N = 3 biological replicates; n = 84 RGCs; *p < 0.05, **p < 0.01, ***p < 0.001; mixed-effects analysis with FDR correction. **c,** Representative images of donor channel (upper row), acceptor channel (middle row), and calculated FRET efficiency of pHLemon-rev in *Vglut2-Cre* pups acquired with 820 nm excitation in the absence of breathing onset. **d,** Quantification showing FRET efficiency in RGCs from newborns without breathing onset after birth. n.s., not significant; N = 5 biological replicates; n = 140 RGCs; mixed-effects analysis with FDR correction. **e,** Summary of changes in intracellular pH, Ca^2+^ levels, ATP levels, AMPK-F activity, and mTOR activity. The results of pHi, SNRK/AMPK-F, Ca^2+^, and ATP are from Fig. 5b (pHi), Fig. 3c (SNRK/AMPK-F), Extended Data Fig. 13d (Ca^2+^), and Extended Data Fig. 13b (ATP). The mTOR results are from Extended Data Fig. 2f.

Because genetically encoded sensors can be pH-sensitive ^64,73,74^, we next asked whether the observed changes in ATP, calcium, or AMPK-F activity could reflect pH-altered sensor properties rather than bona fide signaling. To test this, we expressed ATeam1.03, Twitch-2B, and ExRai-AMPKAR in HEK293T cells and measured their responses across extracellular pH 6.8-7.4 (Extended Data Fig. 18a). Twitch-2B and ATeam1.03 were stable across this range (Extended Data Fig. 18b, c, f, g). ExRai-AMPKAR signal increased as pH rose from 6.8 to 7.4 (Extended Data Fig. 18d, h), consistent with the *in vivo* data, but the phospho-dead mutant sensor did not respond (Extended Data Fig. 18e, i). However, we cannot rule out the possibility that the phosphorylated sensor has a stronger pH dependence compared with the mutant one. Together with the observation of dramatically reduced AMPK-F activation in *Snrk* knockout RGCs (Fig. 3d, e), these results argue that the ExRai-AMPKAR signal detected in RGCs *in vivo* mainly reflects kinase-dependent phosphorylation rather than a direct pH effect on fluorophore behavior.

Together, these data support a two-phase model of postnatal SNRK/AMPK-F activation in RGCs. Early after breathing onset, CO₂ clearance drives rapid intracellular alkalinization without detectable changes in ATP or calcium, consistent with direct pH-dependent activation of SNRK/AMPK-F (Fig. 5e). As calcium subsequently rises, a CaMKK-dependent mechanism may further amplify AMPK-F activity (Fig. 5e). Thus, respiration appears to engage SNRK/AMPK-F through sequential pH- and calcium-linked mechanisms, with early alkalinization providing the most immediate upstream signal for rapid postnatal mTOR suppression.

## CO_2_ inhalation restores mTOR activity and promotes axon regeneration after injury

The findings above point to a candidate mechanism underlying birth-associated mTOR inactivation in RGCs. Given that *Snrk* knockout reactivates mTOR signaling in adult RGCs (Fig. 2), we next asked whether this regulatory pathway remains functionally reversible in maturity. Thus, we induced systemic acidification by exposing adult mice to a CO₂-enriched environment (Fig. 6a). *In vivo* imaging showed that inhalation of 10% CO₂ rapidly lowered intracellular pH in RGCs (Fig. 6b and Extended Data Fig. 19a, b), consistent with previously reported rapid response in pyramidal neurons with CO₂ inhalation ^75^. Intracellular acidification was followed by suppression of SNRK/AMPK-F activity as measured by a decrease in emission ratio from 920/800 nm excitation (Fig. 6c and Extended Data Fig. 20a, b). Notably, these responses occurred on distinct timescales. The CO₂-induced decrease in pHi was rapid, detectable within ∼2 min, and reached a plateau within ∼6 min (Fig. 6b and Extended Data Fig. 19b). By contrast, the reduction in SNRK/AMPK-F activity was delayed, becoming significant only after 1 hr of CO₂ exposure (Fig. 6c and Extended Data Fig. 20b). Thus, extracellular CO₂ appears to trigger a stepwise signaling cascade in which acidification occurs first, followed by attenuating SNRK/AMPK-F signaling. Notably, response kinetics in adult RGCs were slower than those observed during the perinatal period, possibly reflecting developmental maturation of cellular and tissue-level ion/pH homeostatic mechanisms ^76,77^ that occur after birth.

**Figure 6.**
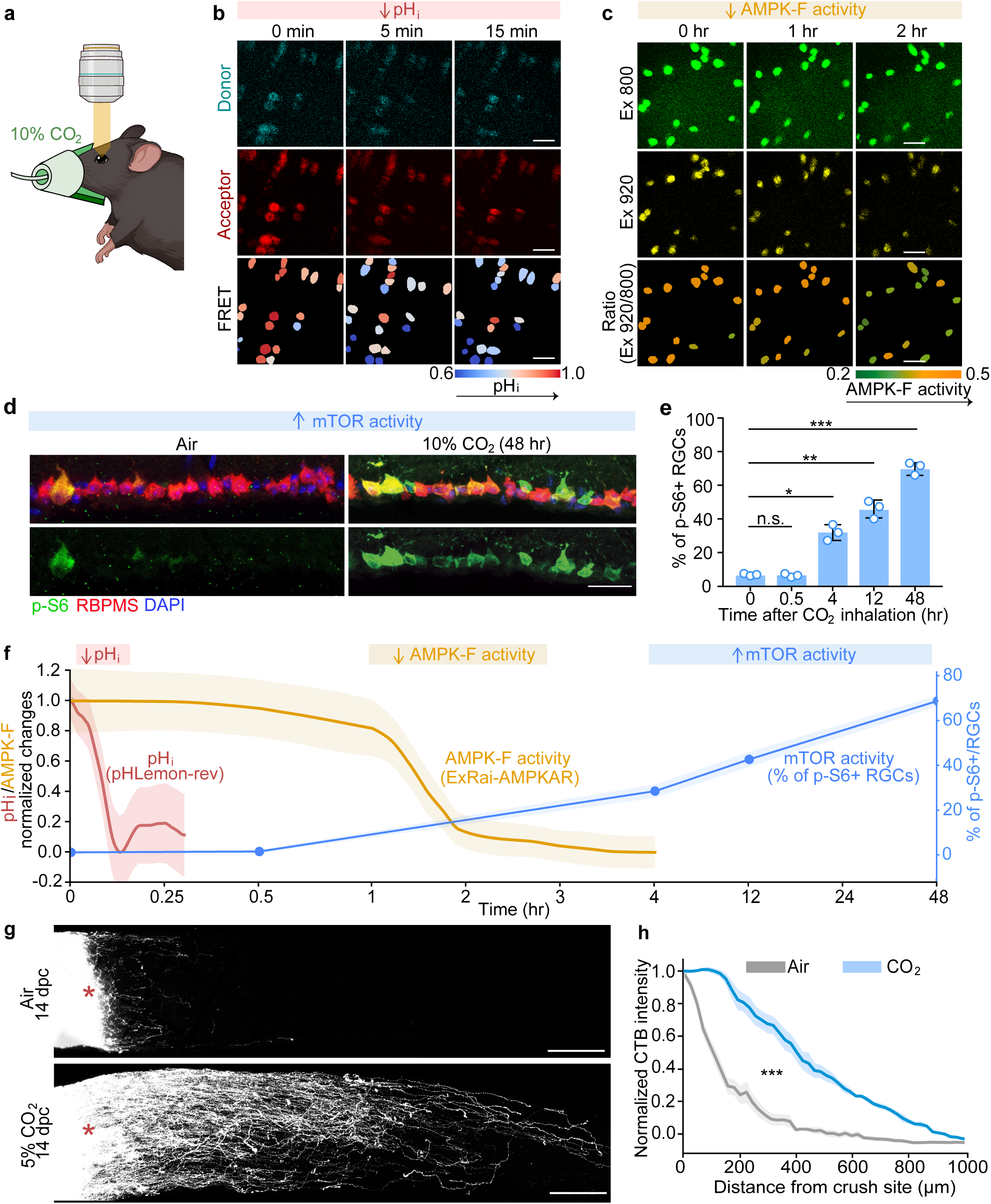
CO_2_ inhalation restores mTOR activity and promotes axon regeneration after injury in adult RGCs. **a,**Experimental scheme of adult mice inhaling 10% CO_2_ for *in vivo* imaging. **b,** Representative images of donor channel (upper row), acceptor channel (middle row), and calculated FRET efficiency (bottom row) of pHLemon-rev in *Vglut2-Cre* adult mice acquired with 820 nm excitation after CO_2_ inhalation at different time points. N = 3 biological replicates; n = 44 RGCs. **c,** Representative images of ExRai-AMPKAR in *Vglut2-Cre* adult mice acquired at 800 nm and 920 nm excitation wavelengths at different time points after CO_2_ inhalation. N = 3 biological replicates; n = 68 RGCs. **d,** Representative images of phosphorylated S6 (p-S6, green), RBPMS (red) staining in retinal sections from adult mice that were primed either in ambient air or 10% CO_2_ for 48 hr. Scale bars, 50 μm. **e,** Quantification showing the percentage of p-S6-positive RGCs at the indicated time points after 10% CO_2_ exposure. Data are presented as the mean percentages of p-S6-positive RGCs among total RGCs in the ganglion cell layer (GCL) of each retina. *p < 0.05, **p < 0.01, ***p < 0.001; n.s., not significant; N = 3 biological replicates for each group; unpaired Welch’s t-test. **f,** Summary of changes in intracellular pH, AMPK-F activity, and mTOR activity. The results of pHi are from Fig. 6b and Extended Data Fig. 19. The results of SNRK/AMPK-F are from Fig. 6c and Extended Data Fig. 20. The mTOR results are from Fig. 6e. **g,** Representative images of optic nerves showing CTB-labeled axons at the lesion site 14 days after injury in wild-type mice. Mice were maintained either in ambient air or in 5% CO₂ starting 7 days before injury and continuing through the 14-day post-injury period. Scale bar, 100 μm. **h,** Quantification of CTB-labeled fluorescent intensity distal to the crush site in wild-type mice that are maintained in ambient air or 5% CO_2_ environment. Data are shown as mean ± SEM; N = 7 and 10 biological replicates for ambient air and 5% CO_2_ conditions, respectively; ***p < 0.001; mixed-effects analysis.

At the downstream end of this pathway, CO₂ inhalation led to a delayed but significant increase in mTOR activity, detectable at 4 hr and reaching ∼70% p-S6-positive RGCs after 48 hr of exposure (Fig. 6d, e). These results support a reversible CO₂/pH–SNRK/AMPK-F–mTOR signaling axis in adult RGCs. The sequential timing of these responses, with pHi after 2 min, SNRK/AMPK-F after 1 hr, and mTOR after 4 hr (Fig. 6f), further argues for a causal relationship between these signaling events rather than parallel independent effects.

Lastly, we tested if driving mTOR activation by CO₂ exposure was sufficient to regenerate injured adult RGC axons. We modified the CO₂ condition for the long term exposure for adult mice. Functionally, this manipulation produced a significant increase in axon regeneration after optic nerve crush (Fig. 6g, h). Together with our finding that conditional deletion of *Snrk* in adult mice restores mTOR activity and enhances axon regeneration after injury (Fig. 2), these data indicate that the regulatory program established during postnatal development remains reversible in the adult. Importantly, in addition to RGCs, CO₂ exposure also reactivates mTOR in the brain, heart and hindlimb skeletal muscle (Extended Data Fig. 21). These results suggested the possibility of a CO₂-based strategy to restore intrinsic regenerative capacity of multiple cell types.

## Discussion

In this study, we identify birth as a critical transition point at which the extrauterine environment reshapes intrinsic cellular states and suppresses regenerative capacity. Specifically, the onset of respiration, particularly through CO_2_ exhalation, induces intracellular alkalinization, activates the AMPK-family kinase SNRK, and suppresses mTOR signaling. Notably, breathing onset-induced mTOR inactivation is not restricted to RGCs, but is also observed across multiple cell and tissue types in the brain, heart and skeletal muscle. Given the conserved and critical role of mTOR in regulating cellular growth and regeneration^4–11,63^, these findings suggest that breathing-associated CO₂ loss may represent a general mechanism for downregulating regenerative capacity after birth. More broadly, our results raise the possibility that the emergence of lung-dependent extrauterine life in amniotes, including mammals and birds, contributes to the markedly restricted regenerative capacity relative to highly regenerative vertebrates such as teleost fish and urodele amphibians^78–80^. These findings suggest that respiratory mechanism and pH regulation may be an underappreciated physiological variable shaping regenerative capacity, providing a testable framework for future comparative studies across vertebrate lineages.

Our findings highlight opposing roles for mTOR and AMPK-family kinase signaling in the regulation of cellular growth capacity. The onset of breathing triggers intracellular adaptations that shift cells from an embryonic, mTOR-driven growth-permissive state to a postnatal, SNRK/AMPK-family-driven state that suppresses anabolic growth and promotes metabolic efficiency. This rapid transition may help postnatal RGCs, and potentially other cell types, adjust to the metabolic and physiological demands of independent extrauterine life, but at the cost of reduced regenerative potential. Although both canonical and non-canonical AMPK-family kinases are expressed in RGCs, our results suggest that SNRK is a major contributor to the AMPK-family activity detected in both perinatal and adult RGCs. Canonical AMPK is generally activated by energetic stress through AMP/ADP sensing by its regulatory γ subunit^47,48^. By contrast, SNRK lacks this canonical regulatory architecture^70^, consistent with our finding that it is relatively insensitive to energy depletion and instead operates through a distinct pH- and calcium-responsive mechanism in RGCs.

Previous studies have implicated altered nutrient availability and oxygen exposure in the birth-associated transition ^44,81^. However, these factors may act on a slower timescale than the rapid mTOR suppression that occurs within minutes after birth. Our results instead suggest that breathing-induced rapid intracellular alkalinization is a key trigger of the switch from mTOR-dominant growth signaling to SNRK/AMPK-family signaling. Both calcium-independent and calcium-dependent mechanisms may contribute to this process, although the underlying details remain to be defined. Intriguingly, this pH-dependent transition resembles sperm capacitation in the female reproductive tract, in which quiescent sperm acquire fertilization competence through bicarbonate uptake and intracellular alkalinization, thereby initiating biochemical and physiological changes, including increased calcium levels, PKA activation, and other downstream events required for fertilization^82,83^. Together, these observations raise the possibility that abrupt shifts in intracellular pH may serve more broadly as efficient triggers of transitions between dormant and active cellular states across diverse biological contexts.

Notably, birth-induced growth suppression appears to be reversible later in life. In adult mice, *Snrk* deletion restores regenerative capacity in injured RGCs after optic nerve crush. In addition, elevated CO_2_ exposure induces intracellular acidification, suppresses SNRK/AMPK-family signaling, and reactivates mTOR in injured adult RGCs, thereby enhancing axon regeneration. Furthermore, CO_2_-induced mTOR reactivation is also observed in other cells in the brain, heart and skeletal muscle (Extended Data Fig. 21), suggesting that this pathway remains responsive to physiological CO₂/pH modulation beyond the retina. Given prior evidence that CO_2_ inhalation has been suggested to suppress seizures in experimental models and patients ^84,85^, our findings raise the possibility of a simple and non-invasive therapeutic strategy that modulates CO_2_ and pH to restore growth and regenerative capacity in RGCs and potentially other cell types throughout the body.

## Methods

### Animals

All procedures performed in this study were in compliance with animal protocols approved by the Institutional Animal Care and Use Committee at Boston Children’s Hospital. Mice were housed in a temperature and humidity-controlled room on a 12-h light/12-h dark cycle (7 a.m. on 7 p.m. off). The animals had free access to food and water. Mice of both sexes were used. Sex was not used as a criterion for inclusion, exclusion or group assignment. Mice of the following strains were used: C57BL/6J (JAX number: 000664), Tsc1^tm1Djk^/J (JAX number: 005680), B6(129S4)-Snrk^tm1.1Hxu^/J (JAX number: 033222), and Slc17a6^tm2(cre)Lowl^/J (JAX number: 016963). We crossed B6(129S4)-Snrk^tm1.1Hxu^/J mice and Tsc1^tm1Djk^/J mice with Slc17a6^tm2(cre)Lowl^/J mice, respectively, to get the conditional knockout mouse strains.

### Plasmids

Plasmids used in this study were obtained from Addgene or generated by standard cloning. Cre-dependent constructs were based on the pAAV-CAG-DIO backbone (Addgene #104052). pAAV-CAG-DIO-pHLemon-reverse was synthesized by Genewiz based on a published sequence ^72^. ExRai-AMPKAR and ExRai-AMPKAR(T/A) inserts (Addgene #192446 and #192447) were subcloned from pcDNA3 vectors into the pAAV-CAG-DIO backbone using the forward primer: 5′-atactttatacgaagttatgctagcggatccgccaccatggtgcggggttctcat-3′ and the reverse primer: 5′-tacattatacgaagttatggcgcgccaagcttttagcgatcaactttgttctgct -3′ (Integrated DNA Technologies). pAAV-CAG-DIO-ATeam1.03 was a kind gift from Philip Williams’s lab at Washington University in St. Louis. pAAV-CAG-Flex-Twitch2B (Addgene #100039), pAAV-hSyn-GFP (Addgene #58867), and pAM-AAV-mSncg-Cre (Addgene #153207) were used as obtained. All constructs were confirmed by Sanger sequencing before use.

### Tissue fixation, sectioning, and immunostaining

Adult mice were transcardially perfused with PBS followed by 4% paraformaldehyde (PFA). Tissues were rapidly dissected and postfixed in 4% PFA overnight at 4 °C. Embryos and timed-delivered newborns were fixed directly in 4% PFA overnight at 4 °C. Fixed tissues were cryoprotected in 30% sucrose overnight at 4 °C, embedded in OCT compound, and frozen at −80 °C. Retina, heart, and hindlimb skeletal muscle samples were cryosectioned at 12 µm and mounted onto Superfrost Plus microscope slides. Brains were cryosectioned at 50 µm and processed as free-floating sections before mounting.

For immunostaining, mounted sections or free-floating brain sections were washed in PBS three times for 10 min each, permeabilized and blocked for 2 h at room temperature in PBS containing 0.1% Triton X-100 and 5% normal donkey serum and incubated with primary antibodies overnight at 4 °C. Sections were then washed in PBS three times for 10 min each and incubated with secondary antibodies for 2 h at room temperature. After final PBS washes, retina, heart, and liver sections were mounted with DAPI Fluoromount-G and sealed. Free-floating brain sections were mounted onto Superfrost Plus microscope slides with DAPI Fluoromount-G and sealed. Sections were stored at 4 °C until imaging. Antibody against phospho-S6 (Ser235/236) is from Cell Signaling Technology (4858S). Antibody against RBPMS is from Novus Biologicals (NBP2-80389).

### Image acquisition

Immunofluorescent images were acquired using laser-scanning confocal microscopes (Zeiss LSM 710, Zeiss LSM 980, or Leica TCS SP8) with 20x and 60x objectives. For quantitative comparisons, images within each experiment were acquired on the same microscope using identical objective, laser power, detector gain/offset, pinhole, scan speed, resolution, and z-stack settings. Images were acquired with software including ZEN Black, ZEN Blue and Leica Application Suite X.

### Quantification and spatial visualization of p-S6-positive RGCs

Retinal cryosections co-stained for RBPMS and p-S6 were used to identify RGCs and quantify mTOR activity at single-cell resolution. For each section, central and peripheral retinal regions spanning 300– 400 µm along the radial axis were analyzed. The central region was selected around the optic nerve head based on DAPI morphology or, when the optic nerve head was not clearly identifiable, positioned symmetrically around the retinal section midline. The peripheral region was selected near the ciliary marginal zone. RGC binary masks were generated from RBPMS images in ImageJ using the IsoData thresholding algorithm, followed by minor manual curation to refine the mask. These RBPMS-derived masks were then used as spatial constraints for semi-automated single-cell segmentation in Python, in which annotated RGC centroids served as seeds for watershed-based segmentation of individual RGCs. Mean p-S6 intensity was quantified within each segmented RGC. The percentage of p-S6-positive RGCs was calculated as the fraction of the segmented RGCs with mean p-S6 intensity above a threshold defined from rapamycin-treated control sections, where p-S6 signal was suppressed to establish baseline-negative levels. Custom analysis scripts are available upon request.

For spatial visualization of p-S6-positive RGC distribution, p-S6 and RBPMS image stacks were manually delineated to the restricted retinal region in ImageJ and imported into Python as 2D projection images. RGC regions were identified from the RBPMS image using smoothing processing, percentile-based thresholding, morphological cleanup, and connected-component filtering. The whole retinal tissue region was similarly defined, and tissue background was estimated from non-RBPMS tissue region. For each channel, the median tissue-background intensity was subtracted, and negative values were set to zero. Spatial aggregation was performed using hexagonal binning implemented in Matplotlib. Within the RBPMS-positive mask, mean background-corrected p-S6 and RBPMS intensities were calculated for each hexagonal bin, and p-S6/RBPMS ratios were computed for bins with detectable RBPMS signal. A tissue-background hexbin layer generated using identical binning parameters was overlaid to preserve retinal anatomical context.

### Immunoblotting

Brain, heart, and hindlimb muscles were rapidly dissected and snap-frozen in liquid nitrogen. For neonatal muscles samples, quadriceps femoris muscles and biceps femoris muscles from both left and right hindlimb were pooled together to generate Western blot sample for each mouse. Tissues were lysed in ice-cold RIPA buffer (Fisher Scientific, PI89900) supplemented with EDTA-free protease inhibitors (Sigma-Aldrich, 11836170001) and phosphatase inhibitors (Roche, 04906845001). Lysates were cleared by centrifugation at 13,000 rpm for 15 min at 4 °C.

Protein concentration was determined using a bicinchoninic acid (BCA) assay (Thermo Scientific, 23227) with bovine serum albumin (BSA; Thermo Scientific, 23209) as the standard. Protein samples were denatured in 4x SDS loading buffer containing 5% 2-mercaptoethanol (Gibco, 21985023) at 95 °C for 10 min, separated by SDS–PAGE, and transferred for immunoblot analysis. Primary antibodies against phospho-S6 (Ser235/236, CST 4858S), phospho-S6K (Thr389, CST 9234T), phospho-4EBP1 (Thr37/46, CST 2855T), total S6 (CST 2217T), S6K (CST 9202S), 4EBP1(CST 9644T) and actin (CST 4970T) were obtained from Cell Signaling Technology. Western blot images were taken using ChemiDoc™ Touch Imaging System (Bio-Rad 1708370). All Western blotting experiments were processed and analyzed using ImageJ version 2.1.0/1.53C.

### Blood glucose measurements

Blood glucose levels were measured using an i-STAT 1 Blood Analyzer (Abbott, 04P75-03) according to the manufacturer’s instructions and previously published protocol ^86,87^ with minor modification. Briefly, ∼90 µL of arterial blood was collected and immediately applied to an i-STAT CG8+ Cartridge (Abbott, 03P88-25). Measurements were performed at room temperature.

### AAV intravitreal injection in adult mice

Intravitreal injection was performed following the previously published protocol ^88,89^. Briefly, mice were anaesthetized with ketamine/xylazine (ketamine 100-120mg/kg and xylazine 10 mg/kg). One drop of Tropicamide was applied to dilate the pupil 10 min before the surgery, followed by one drop of Proparacaine for local anesthesia. Around 1-2 µL of vitreous humor was removed from each eye using a Hamilton syringe and a micropipette. A new micropipette filled with ∼2μL of volume of AAV2 carrying the gene of interest was inserted through the sclera about 0.5 mm posterior to the limbus and into the vitreous chamber without touching the lens. AAV solution (∼2μL) was injected slowly. After injection, antibiotic ophthalmic ointment was applied, and mice were warmed on a heating pad until fully awake.

### Optic nerve crush

Mice were anesthetized with ketamine/xylazine (ketamine 100-120 mg/kg and xylazine 10 mg/kg). We performed optic nerve injury as previously described ^4,88,89^. Briefly, under the microscope, an incision through the dorsal conjunctival membranes was made to expose the optic nerve intra-orbitally. The optic nerve was crushed for ∼1 s, approximately 2 mm behind the optic disc, using trimmed fine forceps (Dumont #5, #11252-00). Antibiotic ointment was applied directly onto the mouse’s eyes to protect the cornea. Animals were closely monitored until full recovery from anesthesia. Mice received analgesic subcutaneously for 24 hours post optic nerve crush.

### Tissue clearing, imaging and quantification of optic nerve regeneration

To visualize regenerated axons, Alexa Fluor™ 555–conjugated cholera toxin subunit B (CTB; Invitrogen, C34776) was injected intravitreally 14 days after optic nerve crush. 48 hours after injection, mice were perfused with 4% paraformaldehyde, and tissues were fixed overnight at 4 °C. Optic nerves were then dissected and cleared using a previously established protocol ^90^ with minor modification. Briefly, nerves were dehydrated through a graded ethanol series (10%, 30%, 50%, 70%, 90% and 100%) for 30 min at each step followed by incubation in Visikol® HISTO-1™ (Sigma, H1-30) overnight at 4 °C. Cleared nerves were subsequently transferred to Visikol® HISTO-2™ (Sigma, H2-30) until imaging.

For imaging, cleared nerves were mounted in Visikol® HISTO-2™ onto Superfrost™ Plus microscope slides. Confocal images were captured in Z-stack scanning mode, and maximum intensity projections were generated to visualize the full extent of labeled axons. Image analysis was performed using ImageJ version 2.1.0/1.53C. The fluorescence intensity profile along the nerve was quantified using the ‘Plot Profile’ function, with background subtraction performed individually for each sample. The crush site was identified on maximum projection images by overlaying five single-plane images acquired at different depths of the same nerve, as previously described ^88^.

### *In vivo* kinase CRISPR screen: sgRNA design and AAV production

A comprehensive list of kinases was assembled using the RIKEN databases ^91,92^. To maximize knockout efficiency, five sgRNAs per kinase were selected from the genome-scale CRISPR/Cas9 knockout (GeCKO) library ^93,94^ to target distinct regions of each gene. Individual sgRNA was subcloned into the PX552 backbone (Addgene 60958 ^95^) under the U6 promoter. Recombinant AAV2 (serotype 2/2) was selected due to its high transduction efficiency and preferential tropism for retinal ganglion cells following intravitreal injection ^4,88,96,97^. All viral vectors were generated by the Boston Children’s Hospital Viral Core. Viral titers used for intravitreal injections were ≥5 × 10¹² genome copies per milliliter (gc/mL) sgRNA construct. For each kinase, the five corresponding sgRNA-expressing AAV vectors were pooled at equimolar ratios and intravitreally injected into adult mice. Each candidate kinase was tested in one animal, with both eyes injected.

To facilitate screening, we established predefined inclusion criteria for selecting hit genes from more than 1,000 kinome and phosphatome targets in the initial screen. Genes meeting these criteria were subsequently validated in independent experiments with 3-4 biological replicates. Specifically, the inclusion criteria are: (i) all RGC axons were fully transected during the crush surgery, and (ii) regenerating axons extended ≥ 500 µm beyond the optic nerve crush (ONC) site. Using these criteria, the forward genetic screen identified 32 hit genes (Extended Data Fig. 3d).

### scRNA-seq analysis

To visualize the developmental expression level of all AMPK-family kinases in RGCs, single-cell sequencing data was retrieved from GEO accession GSE185671 for E13, E14, E16, P0, and P56 RGCs as described previously ^98^. Gene matrices were imported into the R package ‘Seurat’ (version 5.4.0 ^99^) and concatenated into a single Seurat object with all developmental timepoints. Quality control was performed as described previously ^98,100^ to ensure the high quality RGCs transcriptomic profiles. To visualize AMPK-family kinases expression levels, raw counts from biological replicates were averaged using AverageExpression function in Seurat. Expression levels were normalized per gene in Seurat and plotted with the R package ‘pheatmap’ (version 1.0.13 ^101^). To highlight which non-canonical AMPK-family kinases are most highly expressed postnatally, their expression levels were ranked based on mean expression level at the P0 and P56 timepoints and plotted in descending order.

### *In utero* intraocular AAV injection

*In utero* intraocular injection was performed as previously described ^57^. Briefly, pregnant mice with embryos at E13 were anesthetized by isoflurane inhalation. The abdominal skin and wall were cut. The uterus containing embryos was gently pulled out using ring forceps (6 mm OD, FST). Embryos received an intraocular injection of 0.5 μL of AAV2/DJ virus through the uterine wall using glass capillaries (1.0 mm outer diameter; 0.75 mm inner diameter; TW100-4; World Precision Instruments LLC., USA). Micropipettes were pulled with the following setting: HEAT: Ramp+10; PULL: 30; VELOCITY: 120; TIME: 200; PRESSURE: 200 using a Flaming/Brown micropipette puller (Model P-1000; Sutter Instrument Co., USA).

### *In vivo* two-photon imaging setup of retinal ganglion cells

To perform *in vivo* two-photon imaging of RGCs in newborn mice, timed-pregnant mice with embryos at embryonic day 18.5 (E18.5) were anesthetized with ketamine/xylazine (ketamine, 100–120 mg/kg; xylazine, 10 mg/kg). Anesthesia depth was confirmed by loss of toe pinch and palpebral reflexes (∼15 min) and monitored throughout the procedure. Supplemental ketamine (0.066 mg/g body weight) was administered as needed to maintain anesthesia. A laparotomy was performed to expose E18.5 embryos. Embryos were gently removed and positioned in a custom 3D-printed scaffold with the left eye oriented upward. A small incision (∼1 mm) was made in the skin to expose the eye. Optixcare Eye Lub was applied to maintain hydration, and a No. 1 glass coverslip was placed over the eye. Position of embryos was aligned using a rotational stage to ensure that the coverslip was perpendicular to the optical axis. Breathing onset was induced by tactile stimulation using cotton swabs.

To perform *in vivo* two-photon imaging of RGCs in adult mice, mice were anesthetized with ketamine/xylazine (ketamine 100-120 mg/kg and xylazine 10 mg/kg) with their heads fixed to maintain stationary while imaging. A solution of 1% w/v atropine and 2.5% w/v phenylephrine hydrochloride in RO water was applied with an eye dropper for pupil dilation. Mice were placed in the dark for 5-15 min prior to being put under the objective. Optixcare Eye Lub was applied, and the head of the mouse was angled to align the iris with the light path.

RGCs were selectively labeled by intraocular delivery of AAV vectors encoding genetically encoded sensors in the CAG-DIO backbone into *Vglut2-Cre* mice. Viral titers were optimized to 1×10^11^ gc/mL to achieve sparse labeling suitable for single-cell imaging and segmentation. Two-photon imaging was performed using an Olympus FV-MPE-RS multiphoton microscope equipped with a Mitutoyo 20× M Plan Apo NIR objective. Images were acquired at 512 × 512 pixel resolution with 2× digital zoom using galvanometric scanning. The pixel dwell time was set to 2.0 µs, and high-voltage correction was applied. Time-lapse imaging was performed at 1.087 s per frame. Emission signals were collected using three fixed wavelength windows (455-485 nm, 495-540 nm, and 575-645 nm). Before each imaging session, excitation laser power was measured at both the optical bench and under the objective output to ensure consistency.

### Neural Network–Based Cell Segmentation of time-lapse two-photon images

We developed a supervised U-Net-based deep-learning pipeline ^102^ for segmentation of individual cells in time-lapse two-photon images. The dataset comprised 100 manually annotated images spanning multiple experimental conditions. Annotations were generated by three investigators using ImageJ independently. Images were split into training (n = 73), validation (n = 15), and test (n = 12) sets using group-aware partitioning to avoid leakage of similar fields of view. The validation set was used for model selection and hyperparameter tuning, and the test set was reserved for final evaluation.

Images were intensity-normalized, resized to 512 × 512, and single-channel inputs were replicated across three channels for compatibility with the ImageNet-pretrained ResNet-50 encoder ^103,104^. During training, joint image-mask augmentations, including flips, rotations, intensity shifts, and noise were applied using Albumentations ^105^. No augmentation was used for validation or testing. The model used a U-Net architecture with an ImageNet-pretrained ResNet-50 encoder (segmentation_models_pytorch) ^102–104^ and was trained with a composite binary cross-entropy and Dice loss. Optimization was performed with AdamW ^106^ with a two-stage transfer-learning strategy, in which the encoder was first frozen then fine-tuned. Early stopping was based on validation Dice. Hyperparameters, including learning rates, batch size, fine-tuning epochs, frozen epochs, loss weights, augmentation strength, and post-processing thresholds, were optimized using Optuna ^107^. Final model selection was based on validation Dice after threshold optimization.

Model performance was evaluated using Dice, IoU, precision, recall, specificity, and accuracy from pixel-wise predictions after sigmoid activation and thresholding, with small component filtering applied for final evaluation. To assess sample efficiency, models were retrained on increasing dataset sizes. Performance followed a saturating trend, reaching a plateau beyond ∼30 images, and was used to estimate the minimal dataset size required for 95% of maximal performance. The best model was used for subsequent image analysis. Each image was intensity-normalized and resized in the same way as during model training. After binary semantic segmentation based on the optimized threshold, watershed-based post-processing was applied to separate individual cell instances, which were then used for downstream analysis. Custom analysis code is available at https://github.com/HeLabCode/RGC-birth-imaging-pipeline.

### Biosensor selection and sensor-specific imaging strategies

Genetically encoded biosensors were used to monitor intracellular ATP, pH, calcium, and AMPK-family kinase activity in RGCs. AMPK-family kinase activity was measured using ExRai-AMPKAR ^55^. A phospho-null ExRai-AMPKAR(T/A) was used as a negative control ^55^. ATP was measured using ATeam1.03-nD/nA ^65^, intracellular pH was measured using pHLemon-reverse ^72^, calcium was measured using Twitch-2B ^67,68^. These sensors were acquired using two imaging strategies: ATeam1.03, pHLemon-reverse, and Twitch-2B were acquired using single-excitation, dual-emission time-lapse imaging with the frame rate of 1.087s/frame. ExRai-AMPKAR was acquired using dual-excitation, single-emission imaging with sequential alternating excitation at 800 nm and 920 nm across the time course. For ExRai-AMPKAR time-lapse imaging, paired 800 nm and 920nm excitation images were acquired at approximately 1-2 excitation pairs per minute.

### *In vivo* measurement of intracellular AMPK family kinase activity with ExRai-AMPKAR

*In vivo* ExRai-AMPKAR signals were acquired by sequential two-photon excitation at 920 and 800 nm, with emitted fluorescence collected through a common 495-540 nm emission window. To account for the different depth and layer distributions of RGCs during early postnatal stages, z-stack images were acquired for each field of view. Image stacks were collected with an axial step size of 20-30 µm, resulting in about 8-10 optical sections per stack spanning the RGC layer.

Image analysis was performed using a custom Python-based pipeline to process paired stacks of images from 800 nm and 920 nm excitation wavelengths and quantify ratiometric AMPK-F activity. Semi-automated RGC segmentation: Cell segmentation was performed independently for each excitation channel using a semi-automated workflow combining manual seeding and adaptive region growing, guided by a pre-trained neural network as described above. Final cell masks were generated by integrating user-defined seeds with local intensity–based expansion, with neural network predictions used as a reference. In the hybrid mode, region-grown masks were compared to the corresponding neural network–derived instances containing the same seed. Agreement between the two methods was quantified based on fractional overlap, and neural network–derived masks were accepted when overlap exceeded a predefined threshold. In cases of low agreement, segmentation was resolved by selecting either the neural network or region-growing result according to predefined criteria, ensuring robust performance across variable signal and contrast conditions.

Region growing was initiated from user-defined seed points following smoothing of the raw images (Gaussian blur, 3×3 kernel) to reduce pixel-level noise. Seed centers were refined within a local neighborhood by computing the centroid of high-intensity pixels (≥ 80th percentile), providing robust initialization in heterogeneous signals. Local intensity statistics (20th and 80th percentiles, standard deviation) were computed to define adaptive growth thresholds. Expansion criteria incorporated multiple constraints, including a relative intensity drop from the seed, a variance-based term (1.5 × standard deviation), and a local dynamic range term (0.3 × [p80 − p20]), with all terms weighted by local brightness and contrast factors, together with a minimum threshold of 5 intensity units to prevent premature termination in low-contrast regions. The final growth tolerance was defined as the maximum of these constraints. Growth was spatially constrained within a circular window (maximum radius = 25 pixels). This multi-constraint approach enabled consistent segmentation across cells with varying brightness and contrast while minimizing sensitivity to noise and outliers. Segmentation performance was validated against manual annotations across a representative image set spanning different signal intensities, imaging depths, and cellular densities. Comparison between our semi-automatic pipeline and manual annotation was quantified using Dice similarity coefficient, intersection-over-union, centroid displacement, and area error.

#### Z-stack cell tracking

Cell tracking across z-stacks was performed independently for each excitation channel. To correct for axial drift and tissue deformation, three anatomical anchor points were manually selected in each Z-slice and used to estimate slice-wise affine transformations (translation, rotation, and uniform scaling) relative to a reference plane. Following alignment, cell centroids were mapped into a common coordinate system and linked across adjacent z-planes using a nearest-neighbor tracking approach (implemented with Python TrackPy) within a defined spatial tolerance (search range = 20 pixels) and allowing for brief interruptions in detection (up to 2 missing slices). This procedure enabled reconstruction of consistent cell identities across the Z-stack for downstream analysis. To identify corresponding cells imaged under 800 nm and 920 nm excitation, centroid sets were matched in two dimensions after applying a lateral offset to correct residual inter-channel misregistration.

#### Cell pairing across excitation channels

Cell pairing was performed using either global assignment based on the Hungarian algorithm or nearest-neighbor matching with one-to-one constraints. Conflicting assignments were resolved by enforcing uniqueness (each centroid assigned to at most one pair), and unmatched cells were excluded from downstream ratiometric analysis.

#### Illumination-field-based cell exclusion

To account for non-uniform illumination of the retina surface, an illumination mask was computed for each excitation channel independently. Low-frequency illumination fields were estimated by aggregating high-intensity z-slices and applying strong spatial smoothing to define well-illuminated regions. Only cell pairs with centroids located within these regions in both channels were retained for downstream analysis.

#### Ratiometric quantification

For each channel and Z-slice, fluorescence intensity was quantified within segmented cell masks and corrected by subtracting local background estimated from a surrounding annular region that excludes neighboring cells. This accounts for the spatially non-uniform background signals across the imaging field. For each cell, the z-plane with the maximum background-corrected intensity was selected independently for each channel. Ratiometric AMPK-F activity was calculated for each paired cell as the ratio of maximum background-corrected intensities under 920 nm and 800 nm excitation (Ex 920/800). Custom analysis code for all the process described above is also available at https://github.com/HeLabCode/RGC-birth-imaging-pipeline.

#### Optimization of laser power and imaging duration

To determine imaging parameters, laser power was systematically varied from 20% to 70% with 10% interval in our system, resulting in different laser power out of the objective (26 mW, 36 mW, 59 mW, 70 mW, 77 mW, 83 mW with 800 nm excitation wavelength; and 23 mW, 33 mW, 52 mW, 66 mW, 73 mW, 79 mW with 920 nm excitation wavelength) measured by compact power meter console (PM100A, Thorlabs). Increasing laser power resulted in higher fluorescence intensity but also elevated background signal, particularly under 800 nm excitation. A laser power of 40% (59 mW and 52 mW for 800 nm and 920 nm excitation wavelength respectively) was selected as an optimal balance between signal intensity and signal-to-background ratio (Extended Data Fig. 7). Other imaging settings including the PMT voltage, bias voltage, digital gain and digital zoom were kept constant across all images.

To assess photostability and potential photodamage during repeated imaging, sequential excitation at 800 nm and 920 nm was performed for up to 50 cycles with full z-stack imaged at 40% laser power (59 mW with 800 nm excitation and 52 mW with 920 nm excitation). Tissue background intensity was quantified over time as an indicator of photodamage. Based on these measurements, imaging sessions were limited to a maximum of 25 sequential acquisitions to minimize phototoxic effects (Extended Data Fig. 8).

#### Sample size estimation

Quantification was performed at the animal level, with individual cells treated as nested observations using the mixed-effects models. Sample size was determined at the animal level using G*Power ^108^ with an a priori two-tailed independent-samples t-test between two group means. The calculation used an expected effect size cohen’s d = 4, α = 0.05, 80% power, and equal group allocation. This analysis estimated 3 animals per group. Same estimation was done for all *in vivo* imaging data. Quantification was done without sample identity revealed by two independent investigators.

### *In vivo* measurement of intracellular calcium level with Twitch-2B

*In vivo* two-photon imaging for intracellular calcium level with Twitch-2B was performed using single-excitation, dual-emission configuration as described previously ^68^. The Mai Tai pulsed laser was set at

850 nm to most effectively excite the Twitch-2B sensor ^68^. Emission was collected in 455-485 nm (CFP) and 575-645 nm (YFP) wavelengths simultaneously. Cells were segmented using the deep-learning-based segmentation pipeline described above, and background-corrected CFP and YFP emission intensities were averaged within each segmented cell mask at each time point. Twitch-2B signals were quantified as the YFP/CFP emission ratio for each cell, and population-level time courses were generated by averaging per-cell ratios across all analyzed cells at each time point. Raw ratio from time points after breathing onset was normalized to the first frame for visualization.

### *In vivo* measurement of intracellular pH with pHLemon-reverse

pHLemon-reverse consists of a highly pH-sensitive EYFP fused to the relatively pH-stable CFP variant, mTurquoise2. Acidification reduces EYFP fluorescence and increases mTurquoise2 emission by decreasing FRET efficiency. To restrict its expression to the neuronal soma, a C-terminal nuclear exclusion sequence was included. Two-photon excitation was performed at 820 nm. Emitted fluorescence was collected as the donor (455-485 nm; *I_Turq_*) and acceptor (575-645 nm; *I_FRET_*).

We determined the two-photon spectral properties of mTurquoise2 and EYFP using HEK293T cells. Briefly, HEK293T cells were seeded onto glass-bottom imaging dishes (Cellvis, #1 cover glass) in DMEM containing 10% FBS, 100 U mL^−1^ penicillin, 100 μg mL^−1^ streptomycin in a humidified incubator (37 °C, 5% CO_2_). Cells were transfected with mTurquoise2 or EYFP plasmids individually using Lipofectamine ^TM^ 3000 Transfection Reagent (Thermo Fisher L3000001) for 12 hours. Prior to two-photon imaging, culture media were removed, and a #1 glass coverslip was placed over the imaging well without creating air bubbles between the glass coverslip and cells layers. This provided a flat optical interface and reduced the excitation light scattering. HEK293T cells expressing either mTurquoise2 or EYFP were imaged at each wavelength from 720 nm to 960 nm with 20 nm interval. Laser power measured out of the objective to maintain same energy level across spectrum of wavelength. Imaging settings like PMT voltage, bias voltage, and digital zoom were kept constant across all images.

Spectral bleed-through correction was performed to account for both acceptor and donor contamination in the FRET channel, as previously described^69^. Bleed-through coefficients were determined empirically as follows:

#### Acceptor spectral bleed-through (ASBT)

ASBT arises from the direct excitation of EYFP at 820 nm. In cells expressing only EYFP, we first measured the acceptor intensities when excited at both 820 nm and 960 nm to obtain the fraction of acceptor signal directly excited at the donor wavelength. Because the FRET measurements used a single excitation wavelength at 820 nm, we approximated the EYFP intensity excited at 960 nm from the measurement at 820 nm (relative excitation efficiency of EYFP at 820 nm) in cells co-expressing both mTurquoise2 and EYFP. ASBT coefficient (A) was calculated per cell as

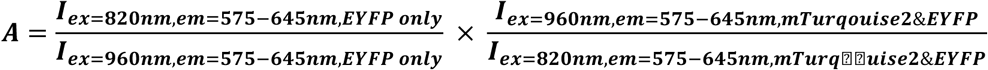

#### Donor spectral bleed-through (DSBT)

DSBT refers to the acceptor channel signal that results from donor emission after being excited at 820 nm. It is determined from cells expressing only mTurquoise2. Its coefficient (B) was calculated per cell as

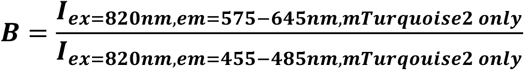

pH dependency of bleed-through coefficients: To determine whether donor and acceptor bleed-through depended on intracellular pH, cells were equilibrated in calibration buffers spanning pH 5.5–8.0 containing 5 µM nigericin (Fisher Scientific, N1495) and 10 µM monensin (Thermo Scientific, 00-4505-51) for 15 min at 37 °C. Bleed-through coefficients did not differ significantly across the tested pH range and were therefore treated as pH-independent constants. Coefficients calculated at each pH condition were averaged across all tested pH conditions, yielding A = 0.016 and B = 0.11.

FRET efficiency calculation: In our system with fixed imaging parameters, with defined bleed-through from donor and acceptor as constants: A = 0.016 and B = 0.11, the subtraction of the SBT signals from the raw acceptor channel (*I_FRET_*) provided the corrected FRET (*FRET_corr_*) as:

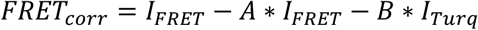

FRET efficiency (*E_FRET_*) was calculated as previously described ^69^ using the following equation:

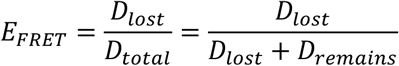

where

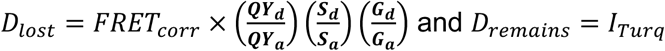

QY refers to the quantum yield, while S and G denote the spectral sensitivity and detector gain of each channel (d = donor, a = acceptor). A fixed quantum yield ratio of 1.474 was used, and 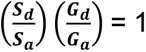 was assumed for all measurements.

Single-cell FRET quantification from *in vivo* time-lapse images: For cell-level quantification from *in vivo* time-lapse images, cells were segmented using the U-net-based network as described above. Pixel-wise FRET efficiencies were averaged within each segmented cell mask to obtain a single mean FRET efficiency per cell at each time point. Population-level time courses were generated by averaging per-cell FRET efficiencies at each time point. Raw FRET values from time points after breathing onset were normalized to the first frame for visualization.

Validation of pHLemon-reverse FRET efficiency calculation: We estimated the intracellular pH using the HEK293T cells transfected with the pAAV-CAG-pHLemon plasmid. Cells were incubated in the calibration buffers (pH range from 5.5 to 8.0) containing 5 µM nigericin (Fisher Scientific, N1495; K*/H* ionophore) and 10 µM monensin (Thermo Scientific, 00-4505-51; Na*/H* ionophore) for 15 min at 37 °C in a humidified incubator (5 % CO,) to equilibrate intra- and extracellular pH. Average FRET efficiencies per cell at different pH levels were computed and fit to a sigmoidal function using L1-norm regularization (Extended Data Fig. 15f):

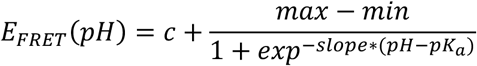

After generating the pH calibration curve and its analytical inverse function:

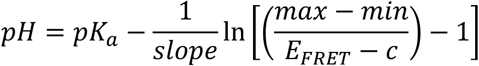

we validated the pH estimation using independent samples equilibrated to known pH values. The pH values estimated from FRET efficiency closely matched the ground-truth pH measured by a pH meter before imaging (Extended Data Fig. 15g).

### *In vivo* measurement of intracellular ATP level with ATeam1.03

ATeam1.03 is a FRET-based ATP sensor composed of CFP and YFP variants linked by the subunit of bacterial ATP synthase, such that ATP binding alters FRET efficiency between donor and acceptor fluorophores. The two-photon spectral properties of CFP and YFP were determined using HEK293T cells as described above. HEK293T cells expressing either CFP or YFP were imaged at each wavelength from 720 nm to 960 nm at 20 nm intervals. Laser power measured out of the objective to maintain same energy level across spectrum of wavelength. Imaging settings like PMT voltage, bias voltage, and digital zoom were kept constant across all images. ATeam1.03 was excited at 820 nm for most efficient CFP excitation, and emitted fluorescence was collected in donor, total fluorescence, and acceptor/FRET channels using 455-485 nm, 495-540 nm, and 575-645 nm detection windows, respectively. Background subtraction, spectral bleed-through correction, cell segmentation, and cell-level FRET-efficiency calculation were performed using the same analysis pipeline described for pHLemon-reverse. Pixel-wise FRET efficiencies were averaged within each segmented cell mask to obtain one mean ATP sensor readout per cell, and population-level traces were generated by averaging per-cell values at each time point. Raw FRET values from time points after breathing onset were normalized to the first frame for visualization.

### HEK293T cell culture

HEK293T cells were obtained from ATCC (CRL-3216). Dulbecco’s modified Eagle’s medium (DMEM), Lipofectamine 3000, Fetal bovine serum (FBS) and penicillin/streptomycin were from Sigma. Adherent mammalian cells were maintained in a humidified 37°C incubator with 5% CO₂ with 10% FBS and 100 IU/ml penicillin, and 100 μg/ml streptomycin and passaged 2-3 times weekly.

### *In vitro* pH sensitivity test of biosensors

HEK293T cells were plated in glass-bottom imaging dishes and transfected with biosensor plasmids around 16 hours before imaging. To modulate the extracellular pH, culture medium was adjusted to defined pH values ranging from 6.8 to 7.4 by HCl and NaOH. Cells were equilibrated for 20 min at 37 °C. Imaging settings were kept constant across the pH series within each experiment.

For analysis, z-stacks were maximum-intensity projected and cells were segmented using Cellpose ^109^. The resulting label masks were used to quantify FRET efficiencies or ratiometric biosensor signals. Background intensity for each channel was calculated from regions outside the cell masks and subtracted from the corresponding channel. Cells with saturated signals were excluded from analysis.

## Acknowledgement

We thank Joshua R. Sanes, Fan Wang and Clifford Woolf for reading the manuscript; Harry Cramer and the Boston Children’s Hospital (BCH) Cellular Imaging Core, funded by NIH grant P50 HD105351 and S10OD03022; Huyan Meng, Lauren Powell and the BCH Viral Core, funded by NIH grants HD018655 and P30EY012196; Yuyang Chen for providing a 3D-printed mouse holder for *in vivo* imaging. The authors used ChatGPT 5.5 to assist in coding and editing the language of this manuscript. This work was supported by the National Institutes of Health (grant numbers R01EY035684 and R01EY036220 to Z.H., R01EY032908 and R01EY035684 to P.R.W., R01MH111647 to H.U., R01GM124038 to G.Y., F32 EY034384 to R.J.D., 5T32EY007145-26 to J.K.K., T32NS007473 to K.C.S.), Gilbert Family Foundation (to Z.H.), Dr. Miriam and Sheldon G. Adelson Medical Research Foundation (to Z.H.), William Randolph Hearst Fund (to X.S.), Goldenson Postdoctoral Research Awards (to X.S.), Herchel Smith Graduate Fellowship (to X.C.)

## Author Contributions

X.S., X.C. and Z.H. conceived and designed the study and wrote the manuscript. X.S. and X.C. collected, processed and performed confocal imaging of samples during perinatal period. X.S., X.C. and D.S. analyzed histology data. X.S., X.C. performed blood glucose measurements. X.S. and M.Y. performed virus injection. X.C. designed plasmid constructs and performed molecular cloning. S.V.H., J.S. and S.M. performed CRISPR-based *in vivo* kinase screen for promoting axon regeneration. X.S., X.C., T.Y., C.Q., P.R.W., G.Y. and Z.H. designed and optimized the *in vivo* imaging system. X.S. and X.C. performed *in vivo* imaging during perinatal and adult stages. X.S., X.C., D.S. and K.C.S. designed and optimized *in vivo* imaging data analysis pipeline. X.S., X.C., D.S. and D.F. processed and analyzed *in vivo* imaging data. D.S. prepared the GitHub repository for codes and pipeline sharing. X.S. and J.Z. optimized the optic nerve surgeries. X.S. and R.J.D. performed the optic nerve surgeries and animal husbandry. X.S. and X.C. and S.Y. performed the biosample collections after surgeries. X.S., X.C., S.Y., and J.C. processed optic nerve tissues. X.S., X.C., S.Y., J.C. and J.K.K. processed retina tissues. J.K.K. performed single-cell RNA sequencing analysis using public a dataset. D.S., R.J.D., J.K.K. and J.C., E.K. contributed to figure illustration and manuscript preparation. All authors contributed to revising the manuscript.

## Competing interests

Zhigang He is an advisor of Axonis and Myrobalan Therapeutics. The remaining authors declare no competing interests.

## Additional information

Correspondence and requests for materials should be addressed to Z.H.

## Code availability

Custom code and pipeline used in this study are available at https://github.com/HeLabCode/RGC-birth-imaging-pipeline. Additional custom scripts are available upon request.

## Data availability

Publicly available database for single cell RNA sequencing analysis in this study can be found in GEO (GSE185671).

